# Genome-Guided Characterization of *Weissella* Species Highlights Strain-Specific Functional Traits Relevant to Food Fermentation and Probiotics

**DOI:** 10.64898/2026.01.23.701356

**Authors:** Nattarika Chaichana, Kamonnut Singkhamanan, Monwadee Wonglapsuwan, Rattanaruji Pomwised, Komwit Surachat

**Affiliations:** Department of Biomedical Sciences and Biomedical Engineering, Faculty of Medicine, Prince of Songkla University, Hat Yai, Songkhla 90110, Thailand; Division of Biological Science, Faculty of Science, Prince of Songkla University, Hat Yai, Songkhla, Thailand

**Keywords:** *Weissella*, comparative genomics, pan-genomeanalysis, exopolysaccharide biosynthesis, probiotic potential

## Abstract

The genus *Weissella* comprises heterofermentative lactic acid bacteria widely distributed across fermented foods, environmental niches, and host-associated habitats, yet the genomic basis underlying their functional diversity and evolutionary plasticity remains incompletely resolved. Here, we performed a large-scale comparative genomic analysis of 347 high-quality genomes representing 16 *Weissella* species to characterize pan-genome architecture, phylogenomic structure, and the distribution of functional traits relevant to fermentation performance, safety, and probiotic potential. Pan-genome modeling revealed a strongly open pan-genome (P = 3419.89⋅N^0.455^), dominated by shell and cloud genes, indicating extensive genomic plasticity and ongoing gene acquisition. Average amino acid identity analysis delineated clear species boundaries while highlighting close evolutionary relationships among several taxa. Functional profiling demonstrated conserved carbohydrate-active enzyme repertoires dominated by glycoside hydrolases and glycosyltransferases, accompanied by species-specific variation in carbohydrate-binding modules. Comparative synteny analysis resolved six major architectures of exopolysaccharide biosynthesis loci, with *W. cibaria* enriched in regulator-rich and potentially active operons. In silico safety screening detected antimicrobial resistance genes in only three strains and no virulence-associated genes across the dataset, supporting a generally low-risk genomic profile. Probiotic-associated traits, including vitamin biosynthesis and stress resistance, were broadly conserved, whereas adhesion, γ-aminobutyric acid biosynthesis, and secondary metabolite gene clusters were rare and strain-specific. Collectively, these results provide a systems-level view of genome diversification within the *Weissella* genus and establish a genome-guided framework for rational strain selection in food, health, and biotechnological applications.

**Importance:** Microbial traits relevant to food fermentation, probiotic performance, and safety often emerge from complex interactions among core metabolism, accessory genes, and genome plasticity, yet these relationships remain poorly resolved at the genus scale. By analyzing 347 high-quality genomes spanning 16 *Weissella* species, this study reveals how an open pan-genome, extensive accessory gene diversity, and lineage-specific gene architectures collectively shape functional potential across this important lactic acid bacterial genus. We demonstrate that key traits, including exopolysaccharide biosynthesis, carbohydrate utilization capacity, stress resilience, and biosynthetic gene cluster distribution, are unevenly structured across species and strains, highlighting the necessity of genome-guided strain selection rather than taxonomic assumptions. The integration of phylogenomic relationships, gene content variation, and functional profiling provides a systems-level framework for linking genome evolution with applied phenotypes. These findings advance our understanding of how microbial genomic diversity underpins ecological adaptation and biotechnological performance and provide a scalable blueprint for rational selection of safe and functionally optimized strains in food and health applications.

## Introduction

The genus *Weissella*, belonging to the family Leuconostocaceae, consists of Gram-positive, catalase-negative, and heterofermentative lactic acid bacteria (LAB). Cells appear as cocci or short rods, typically arranged in pairs or small chains (Teixeira et al., 2021). These microorganisms are widespread in nature and are frequently recovered from fermented foods, plant materials, and the gastrointestinal or oral microbiota of humans and animals, where they contribute to the fermentation of carbohydrates and the production of lactic acid (Fhoula et al., 2022; Fusco et al., 2023). Advances in phylogenetic and genomic studies have refined the taxonomy of *Weissella*. Recent reclassifications clarified species boundaries and introduced the related genus *Periweissella*, providing greater resolution for ecological and biotechnological research within the Leuconostocaceae family. These taxonomic updates are crucial because species identity often determines functional roles, including exopolysaccharide synthesis and probiotic potential, as well as safety considerations (Fanelli et al., 2023). From a technological perspective, *Weissella* species are particularly valued in food fermentation processes. Certain strains are prolific producers of exopolysaccharides (EPS), such as dextran, which can improve the texture, water-binding capacity, and sensory attributes of bread, vegetable ferments, and other products (Hu et al., 2023; B. Zhou et al., 2024). In addition, several strains synthesize bacteriocins, notably weissellicins, that display antimicrobial activity against a variety of spoilage and pathogenic bacteria, making them attractive as natural food biopreservatives (Kim, Yang, & Kim, 2023; Singh et al., 2024). The health-promoting potential of *Weissella* has been investigated most extensively in *W. cibaria* (Liu et al., 2024). Clinical trials, such as a randomized controlled study of *W. cibaria* strain CMU, have demonstrated improvements in periodontal health and modulation of oral microbiota (Do et al., 2025). Laboratory studies of *W. cibaria* strains further support its antimicrobial, anti-biofilm, and immunomodulatory properties against various pathogens, including *Streptococcus pyogenes*, *Staphylococcus aureus*, *S. pneumoniae*, *Moraxella catarrhalis,* and *S. mutans* (Ganguly et al., 2025; Kang, Park, Kim, Lee, & Paik, 2023; Yeu, Lee, Park, Lee, & Kang, 2021). Although promising, these benefits are strongly strain-dependent, and thorough safety evaluations remain essential, particularly as the genus’s broad metabolic capacity can have significant implications for both industrial fermentation processes and human health applications. In this context, our study provides an in-depth comparative genomic analysis of 347 *Weissella* strains, highlighting the functional diversity of these bacteria and offering new insights into their evolutionary trajectories, probiotic potential, and suitability for industrial applications. By integrating genomic perspectives, this research lays the foundation for the rational selection of *Weissella* strains for biotechnological use, emphasizing strain-level validation for safety and functionality.

## Materials and Methods

### 1. Data Collection, Quality Control, and Average Amino Acid Identity (AAI) Analysis

For the data collection of *Weissella* genomes available in the NCBI database, the data retrieval and quality control steps include:

1.1 All 388 *Weissella* genomes were downloaded from the NCBI database (accessed November 3, 2025).

1.2 Quality control (QC) was done to get good-quality genomes by excluding atypical genomes and metagenome-assembled genomes (MAGs). The genomes with >200 contigs were also excluded from the analysis. The quality was assessed by CheckM (Parks, Imelfort, Skennerton, Hugenholtz, & Tyson, 2015), retaining only genomes with estimated completeness ≥95% and contamination ≤5% and Busco completeness ≥95% (Simão, Waterhouse, Ioannidis, Kriventseva, & Zdobnov, 2015).

1.3 Finally, A total of 347 high-quality genomes among the *Weissella* genus were used for further downstream analysis.

1.4 Moreover, all genomes were re-annotated to their taxonomy using the Genome Taxonomy Database Toolkit (GTDB-Tk) (Chaumeil, Mussig, Hugenholtz, & Parks, 2019) and analyzed for average amino acid identity (AAI) among them (Gerhardt et al., 2025).

### 2. Genome Annotation

All the genomes were annotated using the Prokka software using default parameters (Seemann, 2014). The output of the Prokka annotation was used for other identifications.

### 3. Phylogenetic Analysis and Comparative Genomics

The pan-genome analysis of all 347 *Weissella* strains was constructed using the Roary pipeline (Page et al., 2015), where proteins with ≥95% amino acid sequence identity were grouped into the same orthologous family. After running the roary, the pan-genome was divided into the core (*>*99%), soft-core (*<*99% to ≥95%), shell (*<*95% to ≥15%), and cloud (*<*15%) genomes. Pan-genome openness was evaluated using Heap’s law based on the Roary gene presence-absence matrix. Pan-genome accumulation curves were generated from 1000 random permutations of genome order, and the mean pan-genome size was fitted to the model (P = kN^λ^) using log-transformed linear regression, where P is the pangenome size, N is the number of genomes, and k and λ are fitting parameters. The parameter λ was used to assess the openness of the pan-genome. If the value of λ is close to 1, it indicates that the pan-genome is open, meaning it continues to grow as additional genomes are sampled. Conversely, if λ is significantly less than 1, it suggests that the pan-genome size will saturate more quickly and approach a limit, reflecting a closed pan-genome. In addition, a maximum-likelihood phylogenetic tree was generated using FastTree v2.1 (Price, Dehal, & Arkin, 2010) and subsequently annotated and visualized using the Interactive Tree of Life (iTOL) v8.

### 4. In Silico Safety Analysis

Genes potentially linked to safety risks were also predicted. Antimicrobial resistance genes (ARGs) were identified using AMRFinderPlus (NCBI) (Feldgarden et al., 2021) on both the assembled contigs and the Prokka-annotated protein sequences. To independently validate the presence of acquired AMR genes, contigs were additionally analyzed with ABRicate against the ResFinder database, retaining only matches with ≥50% coverage and ≥90% identity (Bortolaia et al., 2020). Putative virulence factors were screened using ABRicate with the VFDB database (Chen et al., 2005), applying more stringent criteria (≥70% coverage and ≥90% identity) to reduce false positives in lactic acid bacteria. ABRicate was executed with default settings following the database setup.

### 5. Genotype Prediction of Carbohydrate-Active Enzymes (CAZymes)

The active genes for the enzymes of carbohydrates in the *Weissella* genomes were identified using dbCAN3 with the DIAMOND tool against the CAZy database (Zheng et al., 2023). The database primarily categorizes enzymes into six major groups, including glycoside hydrolases (GHs), glycosyltransferases (GTs), carbohydrate esterases (CEs), carbohydrate-binding modules (CBMs), auxiliary activity enzymes (AAs), and polysaccharide lyases (PLs).

### 6. Comparative Analysis of EPS Gene Clusters Across Genomes

EPS-related genes were identified in all 347 *Weissella* genomes using the dbCAN3 annotation pipeline (Zheng et al., 2023), with protein sequences analyzed by HMMER to detect CAZyme hits, including GTs and other EPS-associated genes, such as Wzx/Wzy flippases and polymerases. These genes were mapped to their genomic locations using GenBank (.gbk) files, and genomic regions containing co-localized EPS-related genes were extracted as putative EPS biosynthesis clusters. These clusters were grouped into six synteny sub-groups based on their gene order and the presence of key EPS enzymes and regulators. The sub-groups reflect different conserved architectures of EPS loci, with sub-group A representing compact loci with high hypothetical protein content, often seen as truncated EPS segments. Sub-group B contains medium-sized loci with some IS elements. Sub-group C includes full-length loci with conserved EPS genes but low enzyme activity, while sub-group D contains full-length loci enriched with regulators and sugar-modifying enzymes. Sub-group E, the most canonical, consists of full-length loci with enriched GTs, sugar enzymes, and regulators, while sub-group F includes full-length loci with a high presence of IS elements. These cluster-specific GenBank files were compared using clinker (Gilchrist & Chooi, 2021), which generated gene-level homology maps and synteny visualizations for comparative analysis across species.

### 7. Biosynthetic Gene Clusters (BGCs) Analysis

The BGCs associated with secondary metabolites across 347 *Weissella* genomes were detected using antiSMASH v8.0 (Blin et al., 2025), while BAGEL4 was applied to predict gene clusters related to bacteriocins (van Heel et al., 2018).

### 8. Probiotic and Techno-Functional Genes

Probiotic-associated genes related to gastrointestinal (GIT) tolerance, adhesion, biosynthetic gene cluster (BGC)-derived metabolite production, the γ-aminobutyric acid (GABA) system, stress resistance, vitamin biosynthesis, and oxidative stress defense were identified across isolates and linked to species metadata. Gene prevalence was subsequently calculated based on Roary-derived gene presence or absence matrices.

### 9. Data Visualization and Statistical Analysis

All information on species-level strengths was then summarized, and visualization was performed using Python v3.13.7.

## Results

### 1. Genome Distribution Among *Weissella* Species

A total of 388 *Weissella* genomes representing all assembly levels were retrieved from the NCBI RefSeq database on November 3, 2025. To ensure a high-quality dataset for downstream comparative analyses, genome assemblies were curated to confirm accurate taxonomic assignment, assembly quality, and consistent annotation. Genomes classified as atypical assemblies or MAGs, as well as those containing more than 200 contigs or failing basic QC criteria, were excluded. After removing 41 low-quality genomes, a total of 347 high-quality assemblies representing 16 *Weissella* species were retained for further analysis (**Table S1**). The relationship between genome size and GC content across all genomes is illustrated in **Figure 1**, where each species forms a distinct cluster based on its genomic characteristics. The genome sizes of the analyzed *Weissella* species ranged from 1.35 Mbp to 2.49 Mbp, with an average genome length of 1.91 Mbp. The smallest genomes were identified in *W. ceti* (1.35 Mbp), while the largest genome was observed in *W. fermenti* (2.49 Mbp). The GC content varied from 36.58% to 45.50%, with a mean GC content of 41.35% across all genomes. The lowest GC content was found in *W. sagaensis* (36.58%), whereas the highest values were recorded in *W. cibaria*, *W. confusa*, and *Weissella sp.* (45.50%). The species distribution within the dataset was dominated by *W. confusa* (n = 129) and *W. cibaria* (n = 110), followed by *W. paramesenteroides* (n = 56). Moderate representation was observed for *W. viridescens* (n = 12), *W. soli* (n = 8), *W. hellenica* (n = 7), and *W. sagaensis* (n = 6). The remaining species were represented by fewer than five genomes each (**Table S2**).

**Figure 1.**
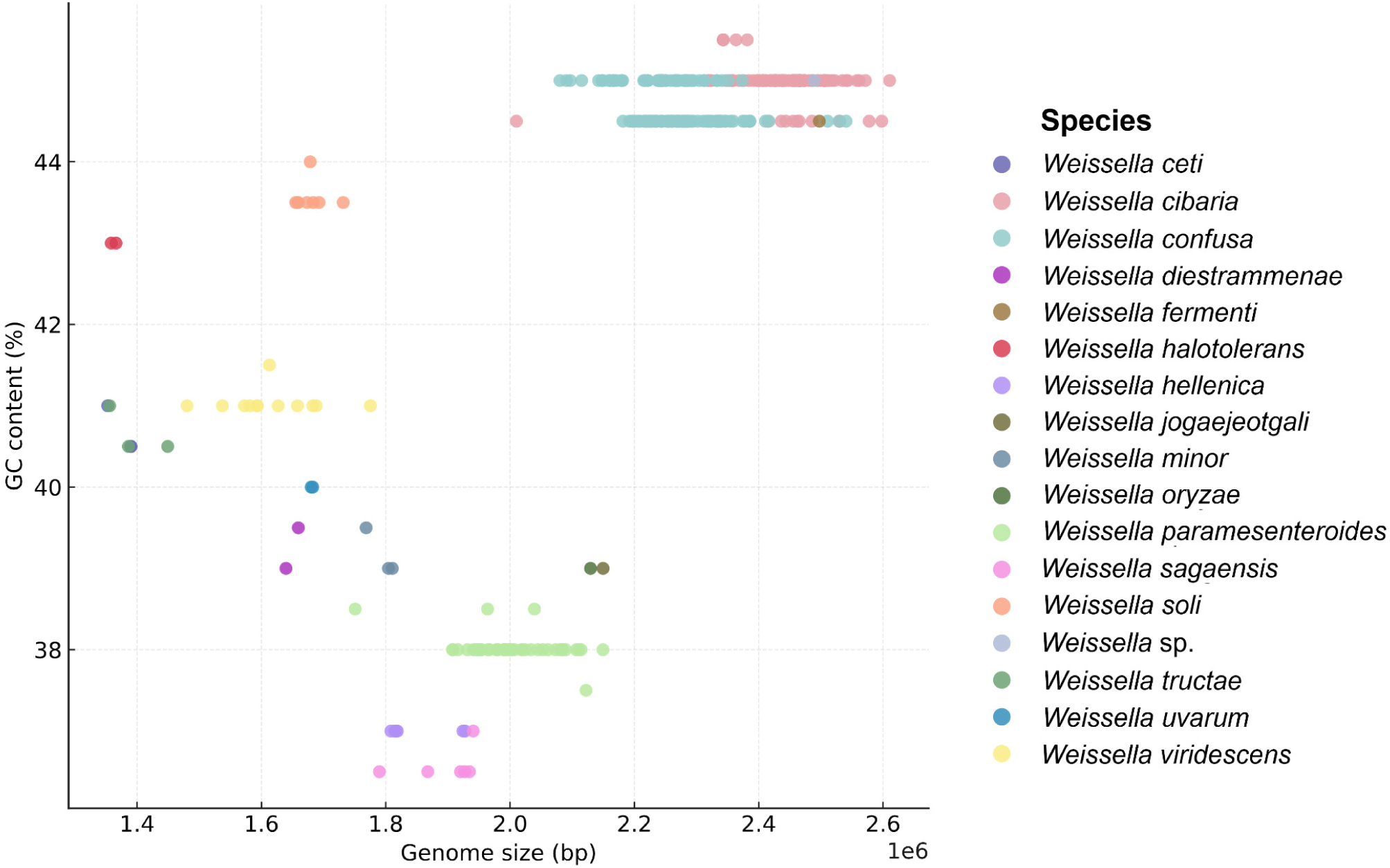
Genome size and GC content of *Weissella* genomes. Scatter plot of 347 *Weissella* genomes showing species-specific clustering patterns across genome size (1.35-2.49 Mbp) and GC content (36.58-45.50%). Each dot represents one genome, colored by species.

### 2. AAI Analysis

The AAI analysis was performed across all 347 high-quality *Weissella* genomes to determine genome-level relatedness and species boundaries. The AAI heatmap revealed distinct, well-defined clusters corresponding to the major *Weissella* species, with clear separation among species-level groups across the dataset. The heatmap showed several distinct diagonal blocks of high AAI values, each corresponding to a group of genomes belonging to the same species (**Figure 2**). The matrices for clusters i to vi contain several small blocks representing species with only a limited number of genomes, including *W. halotolerans, W. uvarum, W. viridescens, W. tructae, W. ceti, W. minor*, and *W. oryzae*. Adjacent to this, a larger block marked as cluster vii corresponds to *W. confusa*, encompassing a large number of genomes and forming a pronounced, continuous high-AI region. Immediately below this, cluster viii represents *W. cibaria*, which forms another large, cohesive, high-identity block occupying a substantial portion of the matrix. Moreover, clusters ix and xii correspond to the group containing *W. hellenica* and *W. soli*, forming a defined rectangular region. Cluster x represents *W. jogaejeotgali*, appearing as a smaller independent matrix. Cluster xi includes *W. paramesenteroides*, forming a distinct high-AI square with moderate size, while cluster 13 corresponds to *W. diestrammenae*, each forming smaller blocks near the bottom of the matrix. Interestingly, *W. tructae* and *W. ceti* are grouped in the same cluster, as are *W. hellenica* and *W. sagaensis*. Other species are distinctly separated into individual clusters, clearly reflecting species-level differentiation.

**Figure 2.**
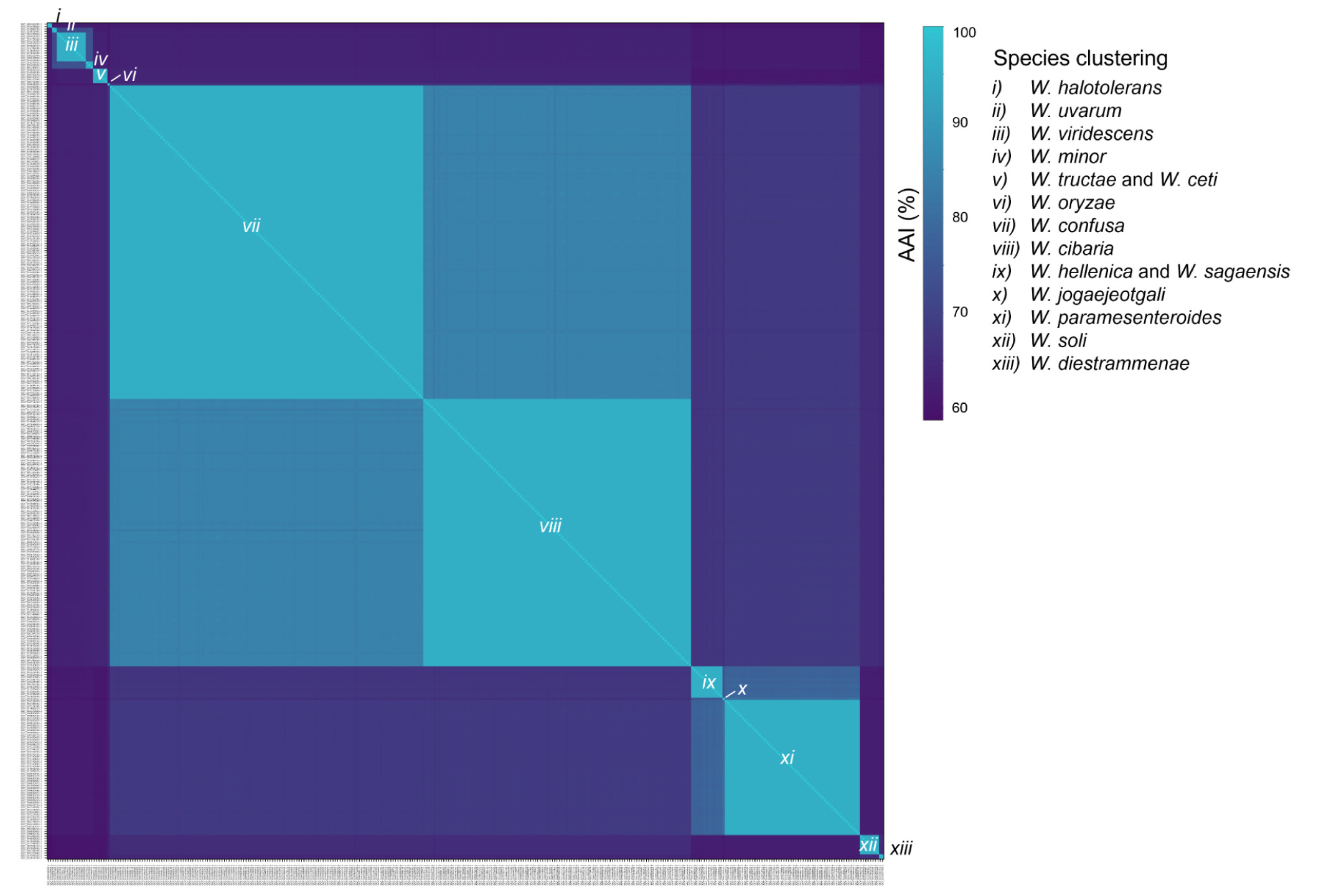
The AAI heatmap of 347 *Weissella* genomes. Heatmap showing pairwise AAI values among 347 *Weissella* genomes. Distinct clusters correspond to major species groups, with clear separation between species. Smaller blocks represent species with fewer genomes, and the lower AAI values between species are visible outside the diagonal clusters.

### 3. Pan Genome Analysis and Phylogenomic Tree Construction

The pan-genome analysis of 347 *Weissella* strains revealed significant genomic diversity, as illustrated by **Figure 3**. A total of 46,741 gene clusters were identified, which were categorized into core, soft core, shell, and cloud genes based on their distribution across the strains. The core genes (7 genes) are present in 99% to 100% of the strains, while the soft-core genes (9 genes) are present in 95% to 99% of the strains. The shell genes (4,965 genes), found in 15% to 95% of the strains, and the cloud genes (41,760 genes), present in less than 15% of the strains (**Figure S1)**. Moreover, Heap’s law fitting of the openness pan-genome followed the power-law model P = 3419.89⋅N^0.455^. The exponent (0.455) indicates that the pan-genome is open, indicating that the pan-genome size continues to increase as additional genomes are included, with no evidence of saturation across the sampled genomes.

**Figure 3.**
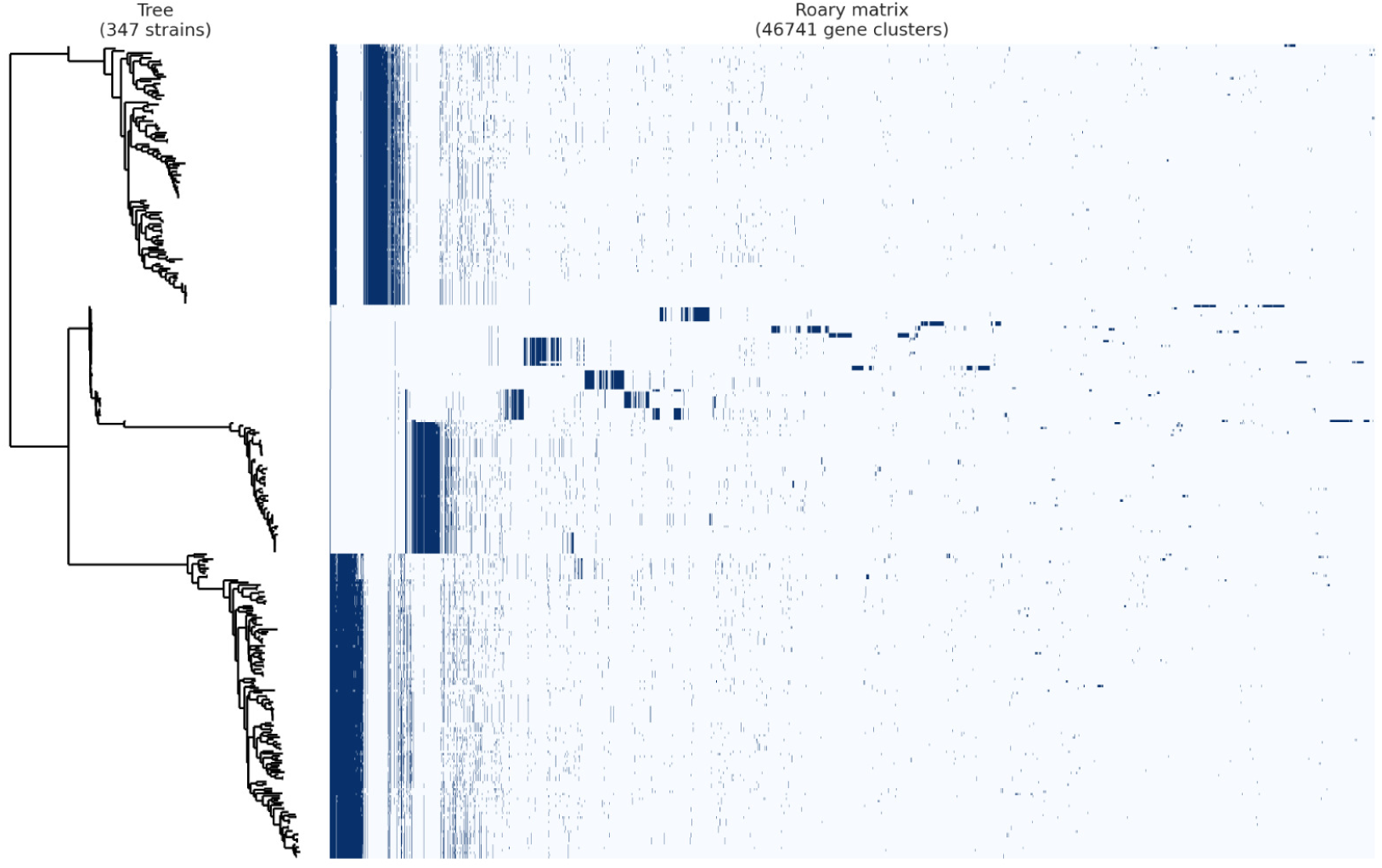
Pan-genome analysis of *Weissella* strains. The pan genome matrix shows the presence or absence of 46,741 gene clusters across 347 *Weissella* strains, categorized into core genes (7 genes, 99%-100% present), soft core genes (9 genes, 95%-99%), shell genes (4,965 genes, 15%-95%), and cloud genes (41,760 genes, 0%-15%). The hierarchical tree on the left represents strain clustering based on genetic similarity, illustrating the high genomic diversity within the genus.

In addition, a phylogenomic tree was constructed using a binary accessory gene presence or absence matrix, representing 46,734 accessory gene clusters across the 347 *Weissella* genomes (**Figure 4**). In this analysis, genomes were grouped based on shared accessory gene content rather than core genomic sequences. Each terminal branch corresponds to an individual strain, and clustering patterns reflect the distribution of accessory genes among genomes. The largest clades also presented species with the highest number of genomes, including *W. confusa* and *W. cibaria*, which appear as densely populated branches. *W. paramesenteroides* forms a separate, cohesive clade, while species with fewer representatives, such as *W. oryzae*, *W. fermenti*, and *W. jogaejeotgali*, appear as small, isolated branches. Several species, including *W. viridescens*, *W. soli*, *W. hellenica*, and *W. sagaensis*, form independent monophyletic clusters with clear separation from other species. However, some species showed accessory genes within other species, such as *W. ceti* and *W. tructae.* Moreover, the 347 *Weissella* genomes were isolated from a wide range of ecological and host-associated environments. These sources encompassed fermented foods, raw food products, animal hosts, insects, human-derived samples, and various environmental niches. The majority of genomes originated from fermented food sources, while additional strains were recovered from animal-derived materials. A subset of genomes was isolated from insect hosts, and a smaller proportion of genomes came from human clinical or commensal samples, as well as environmental sources. *W. confusa* showed the broadest range of isolation sources, including strains from fermented foods, raw food products, environmental samples, and multiple host-associated origins. *W. cibaria* is predominantly represented by strains isolated from fermented foods, and *W. paramesenteroides* includes isolates from fermented foods and animal origin. Species with moderate genome representation, such as *W. viridescens*, *W. soli*, and *W. hellenica*, are isolated from various sources. *W. sagaensis* displays isolates originating mainly from animals. Species with fewer genomes, including *W. ceti*, *W. tructae*, and *W. minor*, consist of strains isolated from fermented foods and animal hosts. Rare species such as *W. oryzae*, *W. jogaejeotgali*, *W. fermenti*, and *W. uvarum* correspond to single-source origins consistent with their individual branches on the tree. Country-of-origin information mapped onto the phylogenomic tree shows the geographic distribution of the 347 *Weissella* genomes across species. Most species originated from multiple countries, forming the broadest geographic representation in the dataset. *W. sagaensis* consists of strains isolated from a specific set of countries, and species represented by fewer genomes, such as *W. ceti, W. tructae*, and *W. minor*, contain isolates from one or two countries. However, rare species, including *W. oryzae*, *W. jogaejeotgali*, *W. fermenti*, and *W. uvarum*, are represented by strains from single country sources corresponding to their individual placements on the tree.

**Figure 4.**
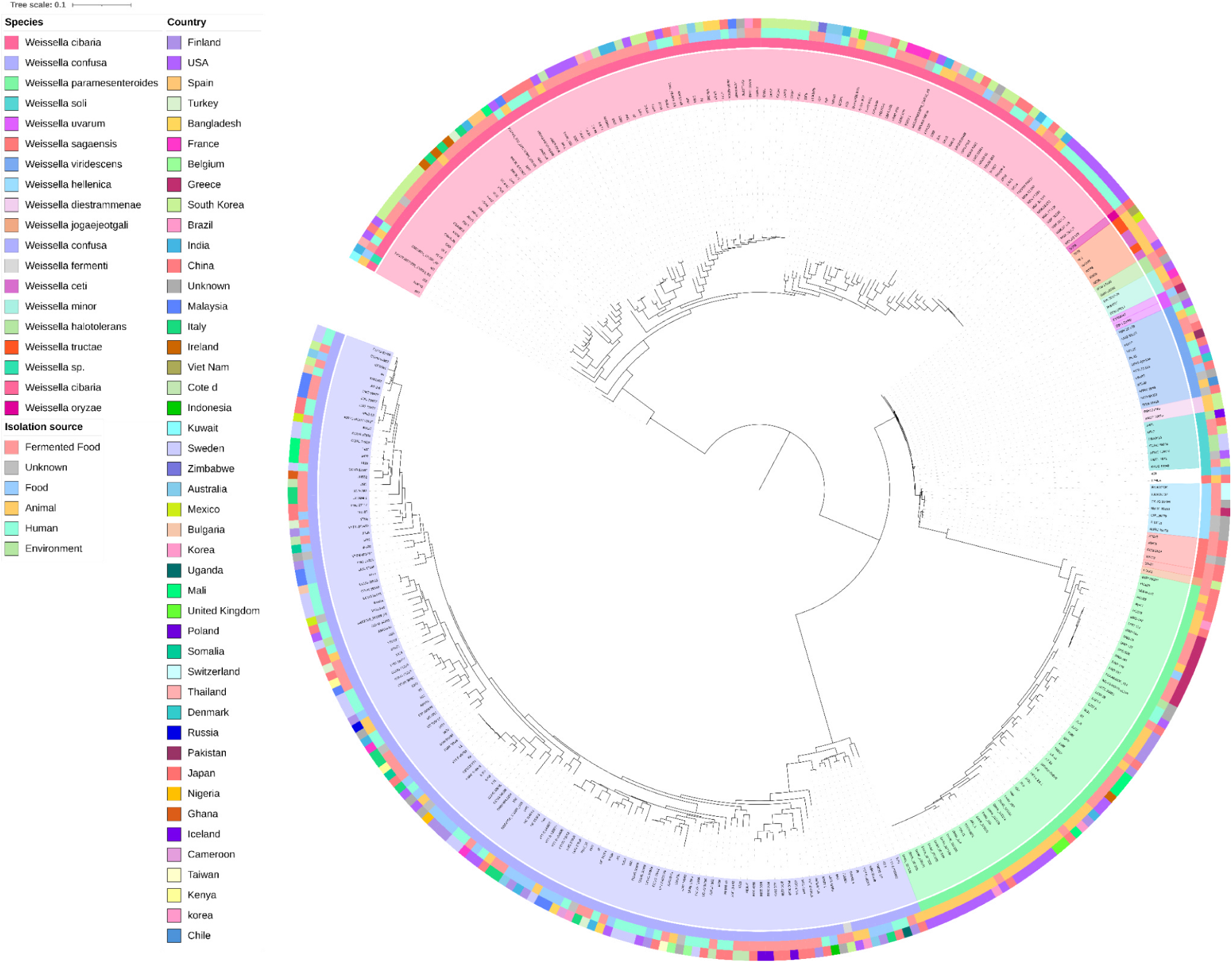
Phylogenomic tree among 347 *Weissella* genomes annotated with species, country of origin, and isolation source. A circular phylogenomic tree was constructed from a binary accessory gene presence or absence matrix to illustrate the genomic relationships among 347 *Weissella* strains representing 17 species. The outer rings display metadata associated with each genome, including the distribution of species alongside their geographic origins and isolation sources across the *Weissella* genus.

### 4. Safety Profile Among *Weissella* Species

Screening of 347 *Weissella* genomes identified ARGs in only three strains, while the remaining 344 genomes contained no detectable ARGs (**Figure 5**). The detected genes spanned several antibiotic classes. The *lnu(D)* gene, present in all three strains, and the *erm(B)* gene, detected in HC968, confer resistance to lincosamides and macrolide-lincosamide-streptogramin B (MLSB) antibiotics, respectively. The aminoglycoside resistance genes *aph(3’)-III*, found in HC330 and HC968, and *ant(6)-Ia*, present in all three strains, are associated with reduced susceptibility to drugs such as kanamycin, neomycin, and streptomycin. The *fexA* gene, identified in HC330 and HC968, encodes resistance to phenicols, while *tet(M)*, detected in all three strains, confers tetracycline resistance via ribosomal protection. The efflux-associated gene *mdt(A)*, observed exclusively in HC970, represents a multidrug efflux determinant linked to decreased susceptibility to multiple antibiotic classes. The genes *poxtA* (HC330 and HC968) and *optrA* (HC330) mediate resistance to oxazolidinones and phenicols, with *poxtA* additionally providing tetracycline resistance. The ionophore-associated genes *narA* and *narB*, detected only in HC330, are linked to resistance to polyether ionophores. No additional ARGs were identified in the remaining genomes, and no virulence factor-associated genes were detected in any of the 347 *Weissella* genomes.

**Figure 5.**
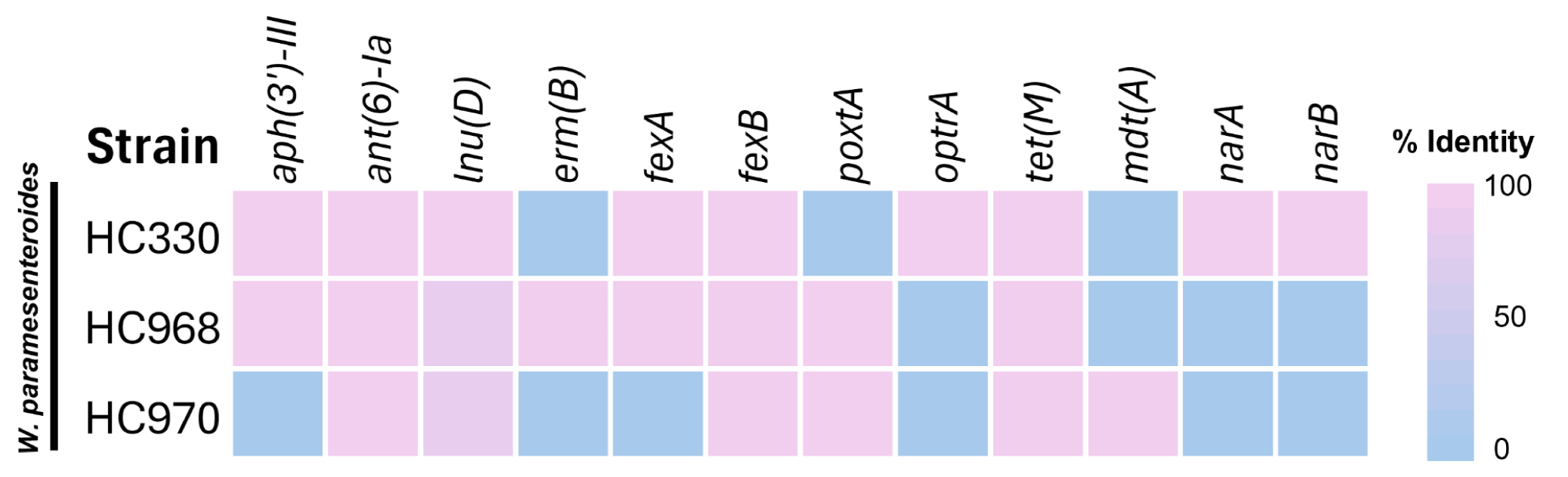
Distribution of ARGs among 347 *Weissella* genomes. Heatmap showing the presence of ARGs across all analyzed *Weissella* strains. Only three strains carried detectable ARGs, while the remaining 344 strains showed no ARGs in the database screen. Each column represents an individual resistance gene, and each row corresponds to a strain, with colored cells indicating gene presence.

### 5. CAZyme Distribution Among *Weissella* Species

The CAZyme analysis revealed a highly conserved distribution among *Weissella* species. In all species examined, GHs constituted the largest proportion of the CAZy repertoire, accounting for approximately 60-70% of the total CAZymes per genome. This dominance was consistently observed across both highly represented species (*W. confusa*, *W. cibaria*, and *W. paramesenteroides*) and moderately represented species. GTs represented the second most abundant CAZy category, contributing roughly 20-30% of the total CAZy content in each species. The relative proportion of GTs was stable across species, with only minor interspecific variation. However, CBMs were present at lower proportions compared with GHs and GTs but showed noticeable variation among species. In particular, *W. confusa* exhibited a higher relative CBM contribution than other species, whereas reduced CBM proportions were observed in *W. paramesenteroides* and several moderately represented species. In contrast, AAs and PLs together accounted for only a small fraction of the CAZy repertoire, typically less than 5% of the total CAZymes. These categories were sporadically distributed and absent in several species, indicating limited representation across the genus (**Figure 6A**). Among these categories, the heatmap analysis of CAZy sub-family prevalence revealed a distinct yet structured distribution pattern across *Weissella* species (**Figure 6B**). A set of core GH sub-families showed consistently high prevalence across all species within the genus. These conserved sub-families were present in both dominant species (*W. confusa, W. cibaria*, and *W. paramesenteroides*) and less represented taxa. In contrast, accessory CAZy sub-families exhibited marked species-specific variation. Several GH sub-families and CBM-associated modules displayed heterogeneous prevalence patterns, with higher representation in *W. confusa* and *W. cibaria* compared with other species. Moreover, *W. paramesenteroides* showed an intermediate distribution pattern, retaining the conserved core sub-families but exhibiting reduced prevalence of accessory components. Moderately represented species, including *W. viridescens, W. soli, W. hellenica*, and *W. sagaensis*, generally showed narrower CAZy sub-family profiles, characterized by the presence of core GH sub-families but limited diversity of accessory sub-families. Several CAZy sub-families were detected only sporadically or at low prevalence, indicating restricted distribution among specific species.

**Figure 6.**
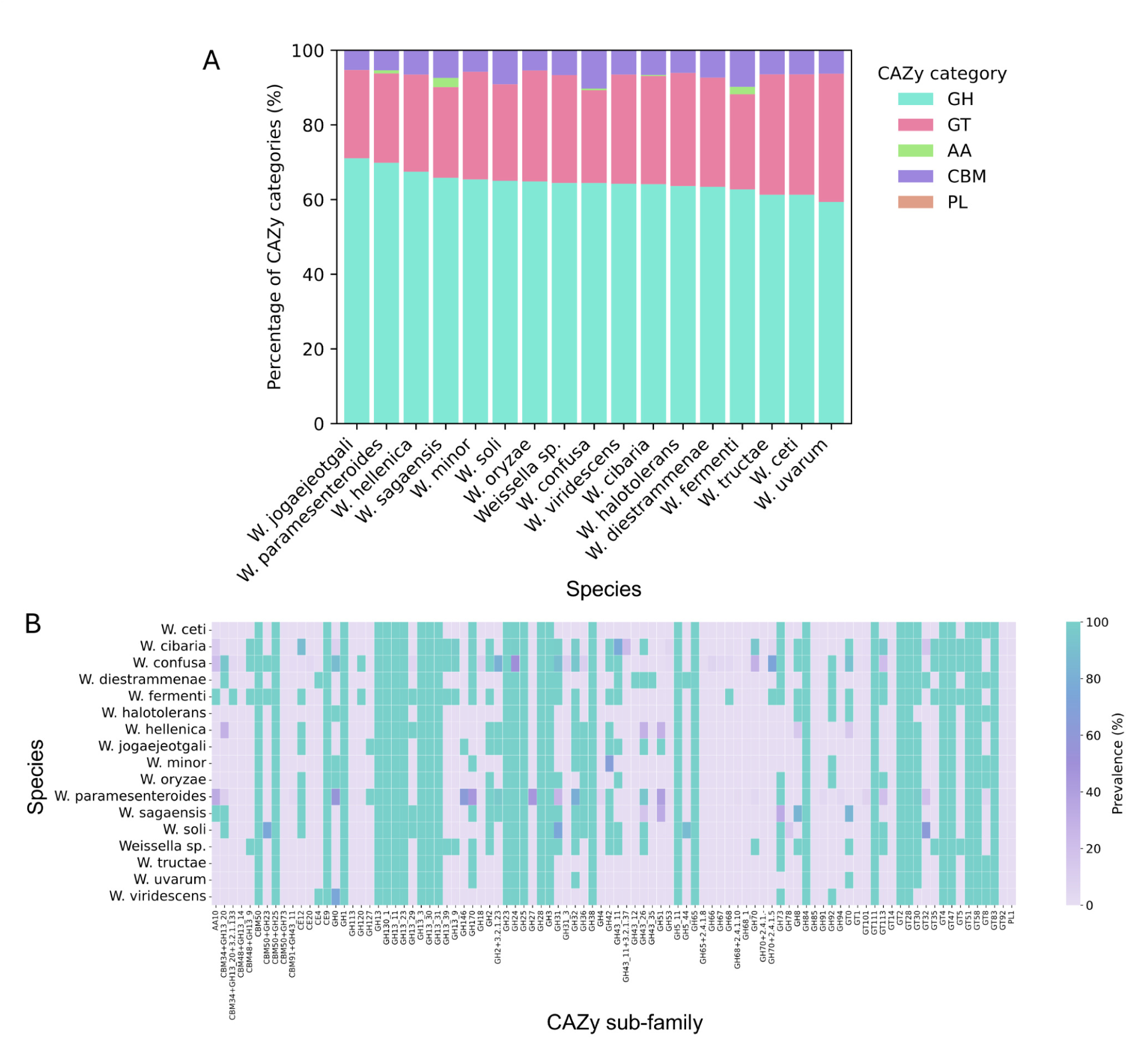
Relative distribution (%) of CAZy categories across *Weissella* species. **(A).** For each species, bars represent the proportional contribution (%) of GH, GT, AA, CBM, and PL, calculated from the mean number of CAZymes per genome, which colors indicate CAZy categories. **Heatmap showing the prevalence (%) of CAZy sub-families across *Weissella* species (B).** Color intensity indicates the prevalence of each sub-family within a species.

### 6. Distribution of Probiotic-Associated Functional Traits Across *Weissella* Species

The distribution of probiotic-associated genes across *Weissella* species revealed a conserved yet heterogeneous functional landscape. Across all species examined, vitamin biosynthesis constituted the largest proportion of probiotic-associated genes, consistently accounting for approximately 45-60% of the total functional repertoire. This dominance was observed across all taxa, indicating that vitamin biosynthesis represents a core probiotic function within the genus. Genes associated with stress resistance, including both general stress resistance and oxidative stress resistance, formed the second most abundant functional group. These traits contributed approximately 30-45% of the total probiotic gene content, with notable enrichment in species such as *W. paramesenteroides*, *W. oryzae*, *W. hellenica*, and *W. soli*. The distribution of GIT tolerance (acid and bile salt tolerance) genes was relatively uniform among species, generally comprising 12-30% of the functional profile. Several species, including *W. halotolerans*, *W. uvarum*, and *W. tructae*, exhibited higher proportions of GIT tolerance genes, while others presented species-specific proportions. In contrast, adhesion- and colonization-related genes were present at low to moderate levels and displayed pronounced interspecies variability. While some species, such as *W. confusa* and *W. cibaria*, showed modest enrichment of adhesion-related traits, many species contained only minimal representations of these functions. Moreover, traits associated with BGCs encoding antimicrobial metabolites and GABA production were rare across the genus (**Figure 7A**).

**Figure 7.**
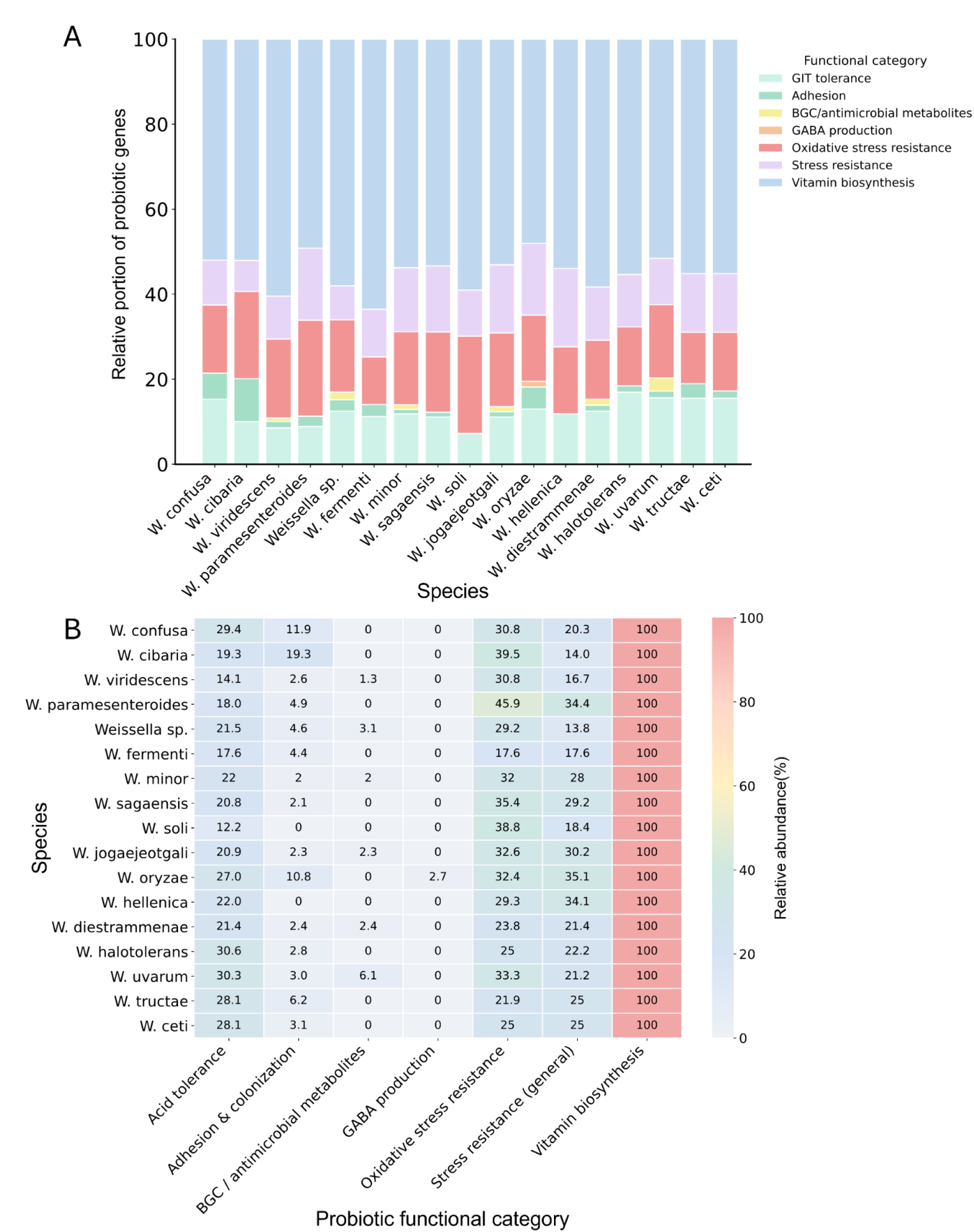
Distribution of probiotic-associated functional traits across *Weissella* species. Relative functional composition normalized to 100% per species (A) and corresponding heatmap of percentage values across functional categories (B).

The heatmap visualization highlights the relative abundance and interspecies variability of functional categories across *Weissella* species. Each value represents the percentage contribution of a given functional category within a species, allowing direct comparison of functional composition. Vitamin biosynthesis is the most dominant functional category across all *Weissella* species, uniformly reaching 100% normalization and representing the largest proportion of probiotic-associated genes. Genes related to oxidative stress resistance and general stress resistance showed moderate to high relative abundance across species, with values typically ranging between 20-45%. Several species, including *W. paramesenteroides*, *W. oryzae*, and *W. hellenica*, exhibited particularly high levels of stress-related traits. Also, GIT tolerance genes are broadly distributed across all species, with relative abundances generally ranging from 12-30% while adhesion and colonization-related genes displayed lower relative abundance and greater interspecies variation. Notably, BGCs encoding antimicrobial metabolites and GABA production genes were either absent or detected at very low relative abundance in most species. Their sporadic distribution across the heatmap underscores their rarity and species-specific occurrence (**Figure 7B**).

### 7. Species-Specific Probiotic Functional Profiles of *Weissella*

Species-specific probiotic functional profiles among *Weissella* species were visualized based on the normalized distribution of probiotic-associated functional categories. *W. confusa* exhibited a relatively balanced functional profile, characterized by strong contributions from vitamin biosynthesis, oxidative stress resistance, and general stress resistance, together with moderate levels of GIT tolerance. Minor but detectable contributions from adhesion-related traits and antimicrobial metabolite-associated genes further suggest functional versatility (**Figure S2A**). Similarly, *W. cibaria* showed a well-distributed functional profile, also dominated by vitamin biosynthesis and oxidative stress resistance, with appreciable contributions from stress resistance and GIT tolerance. Adhesion-related traits were present at low levels, while GABA production and antimicrobial metabolite-associated traits were negligible (**Figure S2B**). *W. paramesenteroides* displayed a profile strongly skewed toward stress resistance and oxidative stress resistance, accompanied by a substantial contribution from vitamin biosynthesis. In contrast, adhesion, BGC, and antimicrobial metabolites, and GABA production were minimally represented (**Figure S2C**). *W. viridescens* exhibited a more restricted functional profile, dominated by vitamin biosynthesis and oxidative stress resistance, with relatively lower contributions from GIT tolerance and general stress resistance, and near absence of adhesion and specialized metabolite-related traits (**Figure S2D**). In addition, *W. hellenica* (**Figure S2E**) and *W. sagaensis* (**Figure S2F**) showed similar patterns, with profiles largely driven by vitamin biosynthesis, oxidative stress resistance, and general stress resistance, while maintaining low proportions of adhesion-related functions and negligible contributions from GABA production and antimicrobial metabolites. *W. ceti* (**Figure S2G**) and W. *tructae* (**Figure S2H**) exhibited elevated contributions from GIT tolerance, alongside strong representation of vitamin biosynthesis and stress resistance traits. In contrast, *W. soli* (**Figure S2I**) and *W. minor* (**Figure S2J**) displayed the narrowest functional repertoires, with profiles dominated by vitamin biosynthesis and stress-related traits, and minimal representation of adhesion, BGC of metabolites, and GABA production, indicating limited accessory probiotic functionality.

### 8. The EPS Biosynthesis Cluster Among *Weissella*

The synteny analysis exhibits the organization of EPS biosynthesis loci across various *Weissella* strains, grouped into six sub-groups of 286 clusters based on their gene order patterns. The representative clusters of each sub-group presented in **Figure 8**. Sub-group A contains medium-sized loci with insertion sequences (IS), found in strains such as *W. confusa*. Sub-group B includes full-length loci with conserved gene order but low EPS-enzyme activity, predominantly in *W. confusa* and some *W. fermenti*. Sub-group C contains full-length loci enriched with sugar-modifying enzymes and regulators, suggesting active EPS biosynthesis, mainly found in *W. cibaria*. Sub-group D represents full-length loci with a high presence of glycosyltransferases and regulators, indicating functional and complete EPS operons, primarily in *W. cibaria*. Sub-group E includes full-length loci with a high presence of insertion sequences and transposases, suggesting ongoing recombination and structural rearrangements. However, sub-group F consists of compact loci with a high proportion of hypothetical proteins and fewer EPS-related enzymes. The summary of each synteny category is presented in **Table S3**.

**Figure 8.**
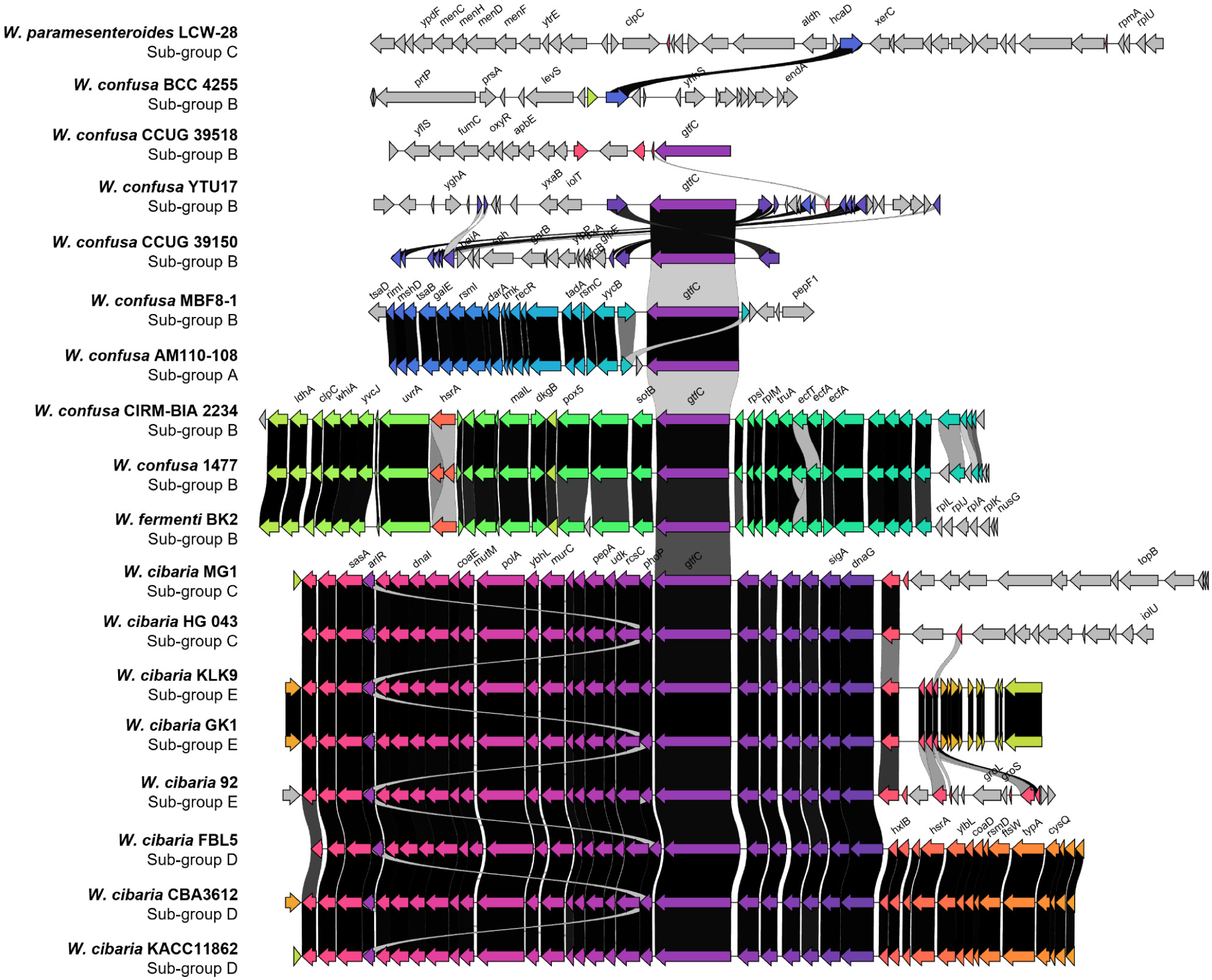
Synteny analysis of representative EPS loci in *Weissella* species. The synteny plot shows six sub-groups of EPS loci based on gene order and functional content. Sub-group A features loci with IS elements, and sub-group B contains conserved loci with few sugar-modifying enzymes. Sub-group C has enriched loci with regulators and sugar-modifying enzymes. Sub-group D shows full-length loci with glycosyltransferases and regulators. Sub-group E is marked by high IS elements and rearrangements, whereas sub-group F consists of compact loci with a high proportion of hypothetical proteins and fewer EPS-related enzymes.

### 9. Prevalence of Biosynthesis Gene Cluster in *Weissella*

The prevalence of secondary metabolite BGCs was analyzed through antiSMASH v.8 across various *Weissella* species. The most common secondary metabolites identified across the species were T3PKS (polyketide synthases) and terpene-precursor clusters, both of which appeared in the majority of species, including *Weissella cibaria* and *Weissella confusa*. Other secondary metabolites, such as RiPP-like, lassopeptides, and lanthipeptide, were found in specific species, such as *W. cibaria* and *W. paramesenteroides*. While these metabolites were less widespread compared to T3PKS and terpene precursors, they were still prevalent within certain species. Interestingly, some species, including *W. viridescens*, *W. confusa*, and *W. paramesenteroides*, exhibited a more restricted repertoire of secondary metabolites. These species exhibited fewer gene clusters, which were often species-specific, such as 2DOS, arylpolyene, and azole-containing RiPPs. These rare metabolites were mostly found in only a few *Weissella* species (**Figure 9**).

**Figure 9.**
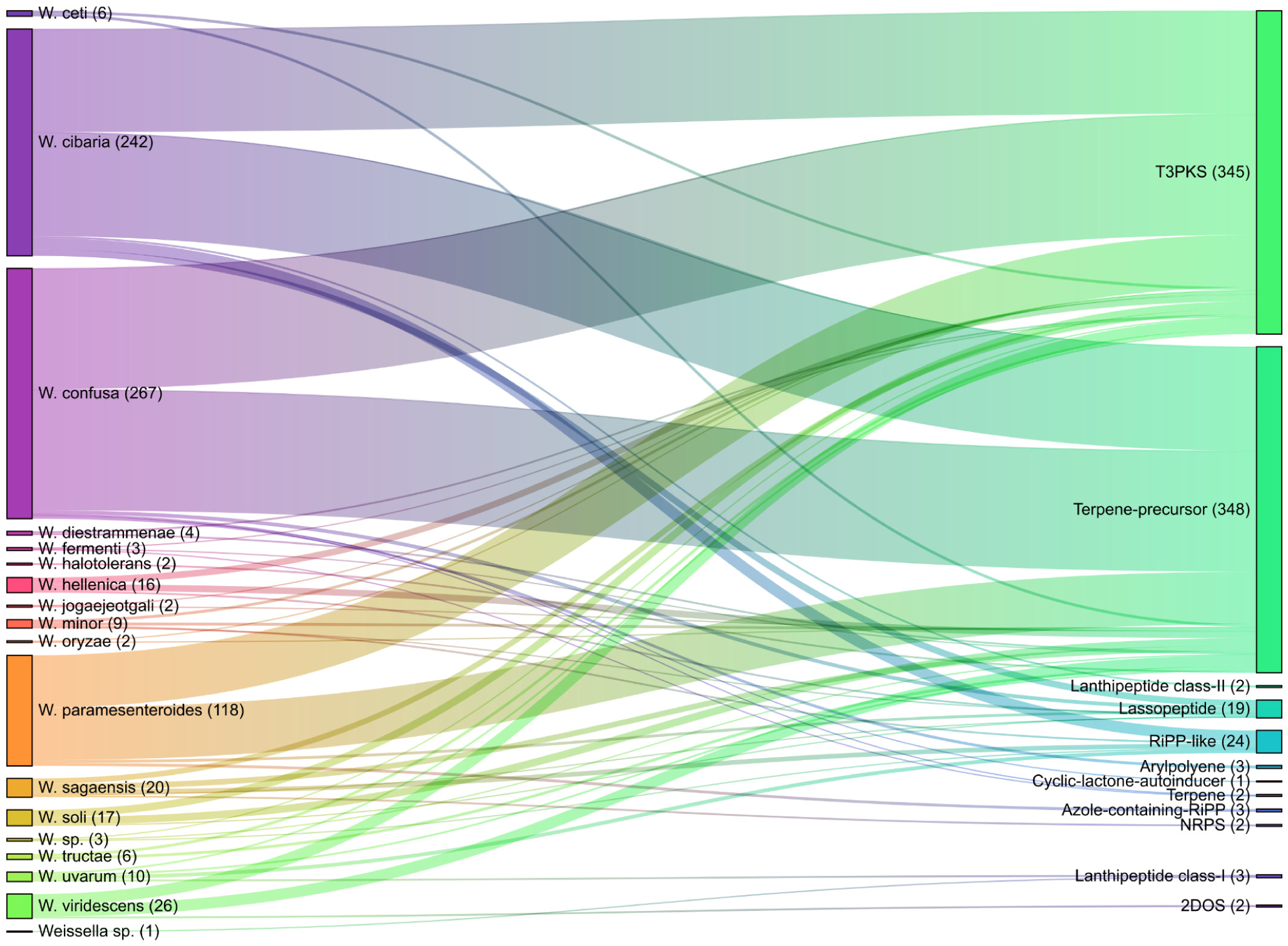
Distribution of secondary metabolite BGCs across *Weissella* species. The sankey diagram illustrates the presence of secondary metabolite BGCs in *Weissella* species. Species are represented on the left, and metabolites on the right, with links showing the presence of gene clusters. Thicker links indicate higher abundance, highlighting core metabolites like T3PKS and terpene-precursor.

## Discussion

The genus *Weissella* comprises heterofermentative LABs widely distributed across fermented foods, plant-associated environments, animal and human-related niches (Sharma, Gupta, & Park, 2023). Comparative genomic studies have shown that *Weissella* species exhibit substantial interspecies variation in genome size, GC content, and accessory gene composition, reflecting evolutionary divergence and niche-specific adaptation. The observed variation in genome size and GC content among *Weissella* species highlights pronounced interspecies genomic differentiation despite a shared phylogenetic background (Apostolakos, Paramithiotis, & Mataragas, 2022). The clear species-specific clustering in genome size-GC content space indicates that these parameters represent stable genomic traits within species, reflecting long-term evolutionary divergence rather than recent horizontal homogenization (Almpanis, Swain, Gatherer, & McEwan, 2018; Ely, 2021). Smaller genomes, such as those observed in *W. ceti*, may reflect genome streamlining associated with niche restriction or host association, whereas larger genomes, exemplified by *W. fermenti*, likely confer greater metabolic and regulatory flexibility (Fahimipour & Gross, 2020; von Meijenfeldt, Hogeweg, & Dutilh, 2023). Similar patterns have been reported in other LABs, where genome expansion is linked to broader ecological adaptability and carbohydrate utilization capacity (Wu, Huang, & Zhou, 2017). The dominance of *W. confusa* and *W. cibaria* in publicly available genomes highlights their ecological prevalence and frequent isolation from fermented foods and diverse environments. In contrast, species represented by fewer genomes showed narrower genomic dispersion, which may reflect limited sampling rather than reduced biological diversity. Nevertheless, the consistent species-specific clustering supports the accuracy of genome curation and validates the dataset as a reliable basis for downstream comparative analyses. For the species identification, the AAI-based clustering revealed sharp species boundaries across the *Weissella* genus, with high within-species similarity and clear separation between species-level groups. The presence of well-defined diagonal blocks corresponding to major species, including *W. confusa*, *W. cibaria*, and *W. paramesenteroides,* confirms the genomic coherence of these taxa. Smaller but distinct clusters representing less abundant species further demonstrate that AAI remains a sensitive and reliable metric for delineating species boundaries even in unevenly represented datasets (Ernster & Rodriguez-R, 2025). Interestingly, the grouping of certain species pairs, such as *W. tructae* with *W. ceti* and *W. hellenica* with *W. sagaensis*, suggests closer evolutionary relationships that may warrant further taxonomic or functional investigation. These relationships are consistent with their proximity in genome architecture and accessory gene content and highlight the value of whole-genome metrics over single-gene phylogenies for resolving fine-scale taxonomic structure within *Weissella* (Abriouel et al., 2015). To translate comparative genomic insights into practical *Weissella* strain selection, a genome-guided, species-informed decision framework was developed (**Figure 10**). This framework integrates safety, metabolic capacity, and techno-functional potential into a sequential selection process while maintaining a clear emphasis on strain-level validation. The first and mandatory selection step focuses on genomic safety screening, where strains are evaluated for the absence of acquired ARGs and virulence factor-associated determinants which are critical consideration when evaluating LABs for food, industrial, and probiotic applications (Haranahalli Nataraj, Behare, Yadav, & Srivastava, 2024). Accordingly, the *in silico* safety assessment of LAB strains represents an important criterion for strain selection and risk evaluation in downstream applications (dR Altavas et al., 2024; Peng, Ed-Dra, & Yue, 2023; Sudheer et al., 2025). The safety screening demonstrated that the vast majority of *Weissella* genomes lack detectable ARGs, reinforcing the general perception of the genus as low risk. The identification of ARGs in only three strains, coupled with the absence of virulence factor-associated genes across all genomes, suggests that resistance acquisition in *Weissella* is rare and strain-specific rather than characteristic of the entire genus. This finding, consistent with the genome-wide analysis, highlights the necessity of excluding non-compliant genomes at the strain level rather than assuming genus-wide safety (Haranahalli Nataraj et al., 2024; Im, Lee, Kim, & Kim, 2023). Following safety clearance, EPS biosynthesis potential represents a key techno-functional decision point. Comparative synteny analysis revealed multiple EPS locus architectures across the genus, allowing species-informed prioritization. Strains of *W. cibaria* are frequently associated with complete EPS loci enriched in glycosyltransferases, regulatory elements, and sugar-modifying enzymes, indicating a higher likelihood of active and structurally diverse EPS production (Y. Zhou, Cui, & Qu, 2019). In contrast, *W. confusa* and *W. paramesenteroides* more commonly harbor compact or conserved EPS loci, suggesting more limited or variable EPS functionality. The variation in EPS loci across species suggests that horizontal gene transfer and mobile genetic elements play significant roles in the evolutionary processes shaping these EPS operons, contributing to the high variability observed across *Weissella* species (Gao et al., 2021). These species-level trends provide an initial filter for applications where EPS-mediated texture or viscosity is a primary objective, while still requiring locus-level confirmation for individual strains.

**Figure 10.**
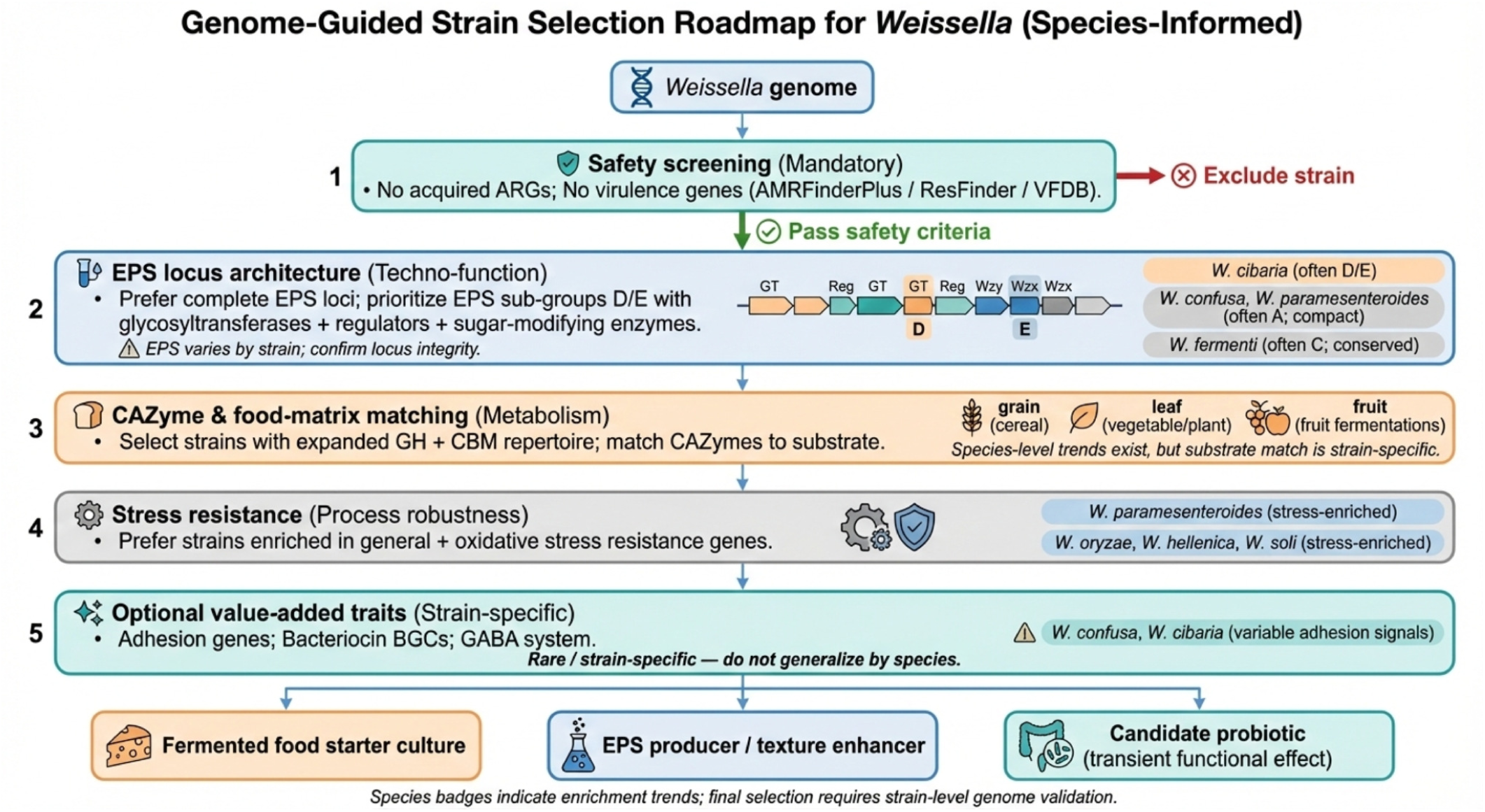
Genome-guided, species-informed decision tree for *Weissella* strain selection. The framework integrates mandatory safety screening with functional genomic criteria, including EPS biosynthesis potential, carbohydrate metabolism, stress resistance, and optional value-added traits. Species labels indicate enrichment trends rather than universal properties, emphasizing the necessity of strain-level genome validation for final selection.

The next decision layer evaluates carbohydrate metabolic capacity, with emphasis on the abundance and diversity of CAZymes-associated genes. Comparative analysis of CAZymes provides key insight into the metabolic capacity and ecological adaptation of *Weissella* species (Liang et al., 2023). The CAZyme analysis revealed a conserved functional backbone across the *Weissella* genus, dominated by GHs and GTs families, reflecting a shared reliance on carbohydrate metabolism. The consistent predominance of GHs across species indicates a core functional requirement for polysaccharide degradation and utilization, particularly relevant to fermented food environments (Paudel, Pardhe, Han, Lee, & Oh, 2025). This finding is consistent with the ecological distribution of *Weissella* in carbohydrate-rich environments, particularly fermented foods and plant-associated niches, where efficient breakdown of complex carbohydrates is essential for growth and survival. The substantial and conserved proportion of GTs suggests that carbohydrate biosynthesis, including cell wall assembly and EPS production, represents another core functional trait of *Weissella*. The relatively higher GT contribution observed in *W. cibaria* may be linked to its documented roles in EPS production and food texture formation, as well as potential interactions with host or environmental surfaces (Wardman & Withers, 2024). In contrast, variation in CBM abundance and CAZy sub-family composition suggests species-specific modulation of carbohydrate-binding and substrate specificity. The enrichment of CBMs in *W. confusa* suggests enhanced ability to bind and process structurally diverse carbohydrates, which may contribute to its ecological versatility and widespread occurrence across diverse fermented food matrices (Olivier, Bull, Bowman, Ross, & Chapman, 2025). The reduction of CBM diversity in *W. paramesenteroides* and moderately represented species may indicate more specialized carbohydrate niches. However, the limited presence of AAs and PLs indicates that *Weissella* species rely minimally on oxidative carbohydrate metabolism and the degradation of uronic acid –containing polysaccharides. This pattern aligns with a predominantly fermentative lifestyle and suggests that these CAZy families represent niche-specific or accessory functions rather than core metabolic traits (Cui et al., 2020). Consequently, CAZyme profiles are best matched to the carbohydrate composition of the target food matrix (e.g., cereal-based, plant-derived, or fruit fermentations), reinforcing a substrate-driven, rather than species-exclusive, selection strategy. The distribution of probiotic-associated traits in *Weissella* species demonstrates a clear distinction between conserved core functions and species-specific accessory traits. Stress resistance and process robustness form an additional selection criterion, particularly relevant for industrial fermentation and product stability. Species such as *W. paramesenteroides*, *W. oryzae*, *W. hellenica*, and *W. soli* show enrichment of genes associated with general and oxidative stress resistance, suggesting enhanced tolerance to acidic, osmotic, and oxidative conditions commonly encountered during fermentation and processing. These traits support their consideration as robust starters or adjunct cultures under challenging production environments (Benhouna et al., 2019; Kaur & Dey, 2023). The strong representation of stress-related genes likely reflects evolutionary adaptation to fluctuating environmental conditions, including acidity, oxidative stress, and nutrient limitation. Species enriched in these traits may therefore be better suited for persistence during food fermentation processes or transient survival in the gastrointestinal tract rather than long-term host colonization. The predominance of vitamin biosynthesis genes suggests that *Weissella* species contribute to micronutrient availability within fermented foods (Keyvan et al., 2025). Conversely, the limited and uneven distribution of adhesion-related genes suggests that host interaction and epithelial attachment are not defining characteristics of the genus as a whole (B. Zhang et al., 2015). Certain species, including *W. confusa*, *W. cibaria*, and *W. oryzae*, exhibited a notably higher presence of adhesion-associated genes compared to others, suggesting that these species possess enhanced adhesive capabilities (Lakra, Domdi, Hanjon, Tilwani, & Arul, 2020; Thant et al., 2024). Optional value-added functional traits, including BGCs and GABA biosynthesis pathways, are incorporated as supplementary selection criteria. These traits play a crucial role in enhancing the functional diversity of *Weissella* species, potentially contributing to probiotic capabilities, fermentation efficiency, and host interactions (Du, Xiong, Xu, Xu, & Wu, 2023). The biosynthetic diversity of *Weissella* species highlights both shared and species-specific pathways for secondary metabolite production. The dominance of T3PKS and terpene precursors across most species suggests that these core metabolic pathways are essential for the survival and adaptation of *Weissella* in diverse environments, such as fermented foods and animal microbiomes. These metabolites are common in many, indicating their role in microbial competition and environmental resilience (Bulnes, Albertos, Jiménez, Castro-Alija, & Díez-Méndez, 2025; Iram, Sansi, Fontana, & Kumar, 2025). In contrast, the presence of RiPP-like, lassopeptides, and lanthipeptides in specific species, such as *W. cibaria* and *W. paramesenteroides,* suggests their ability to produce antimicrobial peptides, which may enhance competitiveness in environments like the gut microbiota or fermented foods (Tenea, Molina, Cuamacas, Marinescu, & Popescu, 2025). The fact that these metabolites are species-specific highlights their potential adaptation to environmental or host-related factors. In contrast, the limited presence of 2DOS, arylpolyene, and azole-containing RiPPs in a few species suggests that these metabolites serve specialized functions, likely related to niche-specific roles such as ecological defense or communication within specific habitats (D. Zhang et al., 2023). These findings emphasize the functional diversity within *Weissella*, where core metabolic functions coexist with a wide range of specialized biosynthetic pathways. This diversity suggests that *Weissella* species are not only adept at fundamental fermentation but also have the capacity to produce bioactive compounds with potential applications in food preservation, probiotic formulations, and biotechnology. However, given their sparse and uneven distribution across the dataset, these traits are considered strain-specific enhancements rather than defining species-level characteristics. This emphasizes the functional plasticity of *Weissella*, where certain strains may possess unique metabolic capabilities that provide ecological or commercial advantages, such as improved gut adhesion, production of bioactive compounds, or stress resistance, while these traits are not necessarily present in all strains of the species. However, the presence of putative probiotic features at the genomic level does not necessarily imply functionality and therefore requires phenotypic validation.

In addition, Heap’s law analysis further supports the conclusion that *Weissella* species possess a strongly open pan-genome. The power-law fit yielded a positive γ-value (0.455) and an α-value below 1 (α = 0.545), indicating that new genes continue to accumulate with increasing genome sampling and that the pan-genome does not approach saturation (Bukhari, Irfan, Ahmad, & Chen, 2022). This quantitative evidence of pan-genome openness is consistent with the overall pan-genome architecture observed in this study, which is dominated by shell and cloud genes and contains only a small core gene set shared across nearly all genomes. The pronounced skew toward accessory genes highlights extensive genomic plasticity within the genus and suggests that *Weissella* species rely heavily on lineage-specific and strain-specific gene repertoires to adapt to diverse ecological conditions (Negrete-Paz, Vázquez-Marrufo, & Vázquez-Garcidueñas, 2025). Such open pan-genome structures are typical of bacteria inhabiting heterogeneous environments, including food fermentation systems, plant-associated niches, and host-associated habitats, where fluctuating selective pressures favor gene gain and functional diversification (Reis & Cunha, 2021). Phylogenomic reconstruction based on accessory gene content further reinforced species-level coherence while revealing substantial functional differentiation within and between species. Major species formed dense and cohesive clades, whereas species represented by fewer genomes appeared as isolated branches, reflecting both ecological specialization and differences in sampling depth (Pontes, Harrison, Rokas, & Gonçalves, 2024). Although occasional overlap in accessory gene content was observed across species boundaries, such as between *W. ceti* and *W. tructae*, these instances were rare and did not compromise overall species separation, suggesting limited historical gene flow or shared ecological pressures. Furthermore, integration of isolation source and geographic metadata demonstrated that pan-genome diversity is closely linked to ecological breadth. *W. confusa*, which displayed the widest range of isolation sources, also exhibited an expansive accessory genome, consistent with its broad ecological distribution. In contrast, species with more restricted ecological niches tended to possess more compact accessory gene repertoires and narrower geographic distributions (Ardalani, Phaneuf, Mohite, Nielsen, & Palsson, 2024; You et al., 2023). These findings indicate that the open pan-genome of *Weissella* reflects ongoing gene acquisition driven by ecological diversity, niche adaptation, and strain-level specialization across environments and regions. Therefore, these findings highlight the genomic flexibility and ecological adaptability of *Weissella*, with significant implications for its application in fermentation, probiotics, and biotechnology, while emphasizing the importance of strain-level validation for accurate functional predictions and optimal strain selection.

## Conclusion

This study provides a comprehensive genomic framework for understanding the diversity and functional potential of *Weissella* species, offering new insights into their ecological adaptation and capacity to produce bioactive compounds relevant to food fermentation, probiotic development, and biotechnology. The strain-level analyses emphasize the importance of genome-guided strain selection, particularly in relation to safety assessment and key functional traits such as exopolysaccharide biosynthesis, carbohydrate metabolism, and stress tolerance, which are critical for performance and stability in industrial applications. The identification of an open pan-genome further highlights the evolutionary plasticity of the genus and its potential as a reservoir for novel functional genes and bioactive metabolites. Therefore, these findings advance our understanding of *Weissella* at both the species and strain levels and support a rational, evidence-based approach to strain selection. This work lays a solid foundation for future investigations into the ecological roles of *Weissella* across diverse environments and for the targeted exploitation of selected strains in functional foods, health-related applications, and sustainable biotechnological innovations.

## Acknowledgements

This research was supported by the National Science, Research, and Innovation Fund (NSRF) through the Program Management Unit for Human Resources and Institutional Development, Research, and Innovation (Grant No. B13F670076). Additionally, funding was provided through the Postdoctoral Fellowship Program from the Faculty of Medicine, Prince of Songkla University.

## Data availability statement

Data will be made available on request.

